# Liver X Receptor regulates Th17 and RORγt^+^ Treg cells by distinct mechanisms

**DOI:** 10.1101/818369

**Authors:** Sara M. Parigi, Srustidhar Das, Annika Frede, Rebeca F. Cardoso, Kumar Parijat Tripathi, Cristian Doñas, Yue O. O. Hu, Per Antonson, Lars Engstrand, Jan-Åke Gustafsson, Eduardo J. Villablanca

**Affiliations:** Immunology and Allergy Unit, Department of Medicine, Solna, Karolinska Institutet and University Hospital, Stockholm, Sweden; Center for Molecular Medicine, 17176 Stockholm, Sweden; Centre for Translational Microbiome Research, Department of Microbiology, Tumor and Cell Biology, Karolinska Institutet, Karolinska Hospital, Stockholm, Sweden; Science for Life Laboratory (SciLifeLab), Stockholm, Sweden; Department of Biosciences and Nutrition, Karolinska Institutet, Stockholm, Sweden; Center for Nuclear Receptors and Cell Signaling, Department of Biology and Biochemistry, University of Houston, Houston, Texas, USA

## Abstract

The gastrointestinal microenvironment, dominated by dietary compounds and the commensal bacteria, is a major driver of intestinal CD4^+^ T helper (Th) cell differentiation. Dietary compounds can be sensed by nuclear receptors (NRs) that consequently exerts pleiotropic effects including immune modulation. However, how NRs regulate distinct intestinal Th subsets remain poorly understood. Here, we found that under homeostatic condition Liver X receptor (LXR), a sensor of cholesterol metabolites, controls RORγt^+^ Treg and Th17 cells in the intestine draining mesenteric lymph node (MLN). Mechanistically, while lack of LXR signaling in CD11c^+^ myeloid cells led to an increase in RORγt^+^ Treg, modulation of MLN Th17 was independent of LXR signaling in either immune or epithelial cells. Of note, LXRα modulated only the Th17 cells, but not RORγt^+^ Treg in the MLN and horizontal transfer of microbiota between LXRα^−/−^ and WT mice was sufficient to partially increase the MLN Th17 in WT mice. While LXRα deficiency increased the abundance of Ruminococcaceae and Lachnospiraceae bacterial families compared to the WT littermates, microbiota ablation including ablation of SFB was not sufficient to dampen LXRα-mediated expansion of MLN Th17. Altogether, our results suggest that LXR modulates RORγt^+^ Treg and Th17 cells in the MLN through distinct mechanisms.

## Introduction

The intestinal barrier is continuously exposed to the changing external environment. To face this challenge, our immune system has evolved to rapidly adapt to environmental changes by mounting tolerogenic and effector responses on demand. A dynamic equilibrium between these two arms of the immune system is required to maintain homeostasis, and failure of mounting an appropriate regulatory response may lead to uncontrolled inflammatory reactions and chronic immune disorders.

Owing to their ability to react in an antigen-specific manner CD4^+^ T helper (Th) cells are central players in ensuring a protective response while limiting collateral damage. Mounting a functional Th cell response in the intestine relies on the anatomical distribution of T cells in immunological inductive and effector sites, where the former is constituted by intestine-draining lymphoid structures such as MLN and the latter being the intraepithelial and lamina propria compartments ^1^. The inductive sites represent the arena where naïve CD4^+^ T cells encounter their cognate antigens for the first time and functionally commit to specific effector subsets. Under steady state, retinoic acid-related orphan receptor γt (RORγt)^+^ Th cells (Th17) and Foxp3 expressing regulatory T cells (Treg) are the most represented Th subsets in the MLN. Despite a certain degree of lineage and functional flexibility, Th17 and Treg cells play complementary roles in maintaining intestinal homeostasis. Although the expression of RORγt and Foxp3 was originally thought to be mutually exclusive, a distinct new population of RORγt^+^Foxp3^+^ regulatory T cells (RORγt^+^ Treg) was described in the mouse intestine ^2-4^, that were shown to control immunosuppression and Th2 responses at the intestinal barrier ^2-4^.

Differentiation to Th17 cells in lymphoid organs is contingent on the presence of TGF-β1, IL-6 and IL-1β cytokines, while their effector function and pathogenicity relies on IL-23. In agreement with their high potential to induce Th17 cell differentiation, intestinal dendritic cells (DCs) are a major source of IL-23, IL-6 and IL-1β ^5,6^. Of note, mice lacking a specific DC subtype characterized by the expression of CD11b and CD103 have reduced numbers of Th17 cells ^6^, highlighting the important role of myeloid cells in Th17 cell homeostasis. The intestinal tract represents a hub for the generation of Th17 cells, which in turn have been shown to exert their effector functions both in the intestine and in extra-intestinal sites ^7^. This phenomenon can be explained by the contribution of environmental signals, mainly diet- and bacterial-derived, in Th17 cell generation. As examples, diets rich in long-chain fatty acids or high in salt content supports Th17 cell differentiation in the gut while germ-free animals have reduced numbers of Th17 cells ^8-10^. Of note, one specific Gram-positive bacterium related to *Clostridia*, segmented filamentous bacteria (SFB), has proven sufficient to induce Th17 cell development in the intestine ^10^. By adhering to the intestinal epithelium, SFB drives the production of serum amyloid A, which in turn boosts the expression of IL-6 and IL-23 by antigen presenting cells, ultimately leading to SFB-specific and unspecific Th17 cell generation in the gut ^10^.

At the crossroads between host and environment, NRs are specialized sensors capable of detecting and functionally translating environmental cues by regulating gene transcription. Liver X Receptor (LXR), a member of the NR family, binds to cholesterol metabolites oxysterols and acts as a master regulator of cholesterol homeostasis ^11^. There are two isoforms of LXR, with LXRβ being ubiquitously expressed and LXRα having tissue specific expression, such as in the liver, adipose tissue, intestine and macrophages ^11^. Both isoforms are expressed and have an antiproliferative effect in human CD4^+^ T cells ^12^, implicating a role of LXR in modulating T cells function. Administration of synthetic LXR ligand *in vitro* and *in vivo* has been shown to promote Treg cell generation while dampening effector T cell responses ^13^. However, other studies have shown no effect in Treg differentiation upon LXR activation *in vitro* ^14^. Importantly, activation of both LXR isoforms plays a part in inhibiting Th17 cell generation in a T cell-intrinsic fashion by inducing the expression of Srebp-1, which in turn physically interacts with and inhibits aryl hydrocarbon receptor (Ahr), a known inducer of Th17 cells ^15^. Consequently, systemic activation of LXR ameliorates symptoms of experimental autoimmune encephalomyelitis (EAE), a Th17-driven mouse model of multiple sclerosis ^15^. While these findings support the notion of LXR as a regulator of Th17 cell biology, whether LXR modulates intestinal T cell homeostasis *in vivo* is yet to be investigated. Here, we investigated the relative contribution of different LXR isoforms in controlling Th17 cells under homeostatic conditions. We found that while both LXR isoforms constrain Th17 cell expansion in the MLN and that LXRβ, but not LXRα, regulates RORγt^+^ Treg homeostasis through LXR signaling in CD11c^+^ myeloid cells. Bone marrow chimera and conditionally deficient mouse models show that this constraint on Th17 cells is independent of LXR signaling not only in T cells, but also in immune and intestinal epithelial cells. Further, while microbiota analysis of LXRα^−/−^ mice, which selectively modulated only Th17 cells, showed increased abundance of bacteria belonging to the family Ruminococcaceae and Lachnospiraceae, microbiota ablation including ablation of SFB was not sufficient to dampen expansion of Th17 in the MLN of LXRα^−/−^ mice. Overall, our study points towards distinct modes of regulation of RORγt^+^ T cells by LXR in the MLN.

## Results

### LXR regulates RORγt^+^ T cell subsets in the MLN in steady state

To evaluate the role of LXR in regulating intestinal Th17 cells, C57BL/6J wild type (WT) mice were administered a modified diet containing GW3965, a well-characterized LXR agonist ^16^. After 10 days, T cell subset composition in MLN was assessed by flow cytometry. While the proportion of CD3^+^CD4^+^ T cells out of total immune cells was unchanged (**Fig. 1a**), lineage specification into RORγt^+^Foxp3^−^ (referred as Th17) and RORγt^+^Foxp3^+^ (referred as RORγt^+^ Treg) cell frequencies were reduced by GW3965 administration compared to control diet (**Fig. 1b and 1c**). These results suggest that activation of LXR *in vivo* restrains T cell differentiation to RORγt^+^ Treg and Th17 cells in the MLN. To further complement these findings, we analyzed the MLN of mice lacking either LXRα (LXRα^−/−^) or LXRβ (LXRβ^−/−^) compared to C57BL/6J WT mice obtained from the same provider. In agreement, deficiency of either isoform of LXR resulted in increased proportion of Th17 cells in the MLN (**Fig. 1d and e**). In contrast, RORγt^+^ Tregs were increased only in LXRβ-deficient animals compared to both WT and LXRα^−/−^ mice (**Fig. 1d and e**). Thus, while LXRβ-deficiency results in expansion of both RORγt^+^ Treg and Th17 cells, LXRα-deficiency resulted in specific expansion of Th17 cells. To gain further insight on how genetic deficiency in LXRα isoform drives proportional expansion of MLN Th17 cells, we tested whether proliferation (Ki-67 staining) or survival (AnnV staining) of Th17 cells was favored in LXRα^−/−^ mice. Although we did not see differences in Ki67^+^ Th17 cells (**Fig. 1f**), we observed a drop in AnnV^+^ Th17 cells in LXRα^−/−^ mice (**Fig. 1g**), suggesting that Th17 cells in LXRα^−/−^ mice have a better survival compared to WT mice. Overall, we showed that while LXR activation regulates the levels of RORγt^+^ T cells in the MLN, we observed an isoform specific effect of LXRβ in modulating RORγt^+^ Tregs and the ability of LXRα in modulating only the Th17 cells but not RORγt^+^ Tregs.

**Figure 1.**
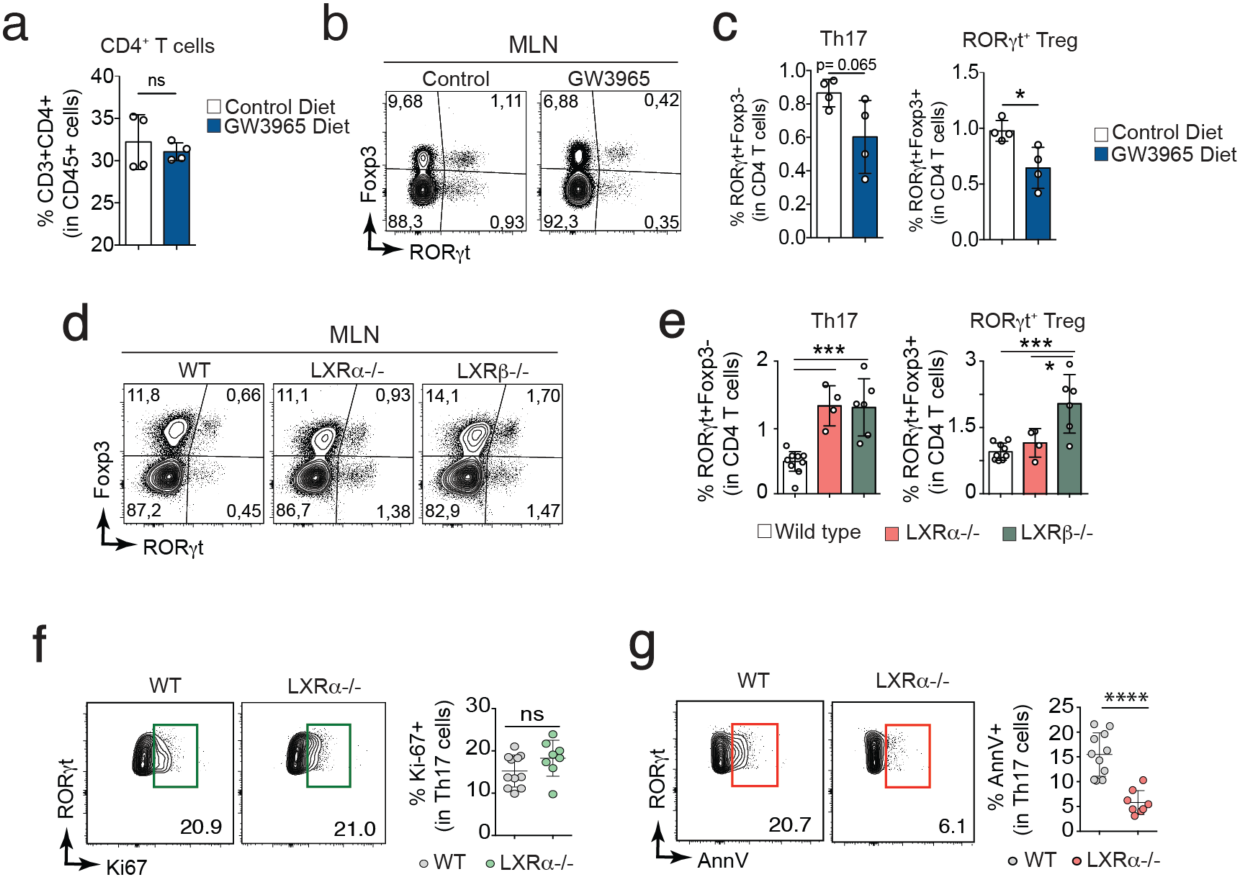
LXR regulates RORγt^+^ T cells in the MLN. **(a-c)** Wild type (WT) mice were fed either a control or GW3965 containing diet for 10 days followed by analysis of T cells in the MLN. **(a)** Frequencies of CD3^+^ CD4^+^ T cells out of CD45^+^ cells in the MLN. (**b-c**) Representative dot plots **(b)** and frequencies **(c)** of RORγt^+^Foxp3^−^ (Th17) and RORγt^+^Foxp3^+^ (RORγt^+^ Treg) cells out of total CD4^+^ T cells (*n*=4 mice for each group). **(d-e)** Representative dot plots (**d**) and frequencies (**e**) of Th17 and RORγt^+^ Treg out of total CD4^+^ T cells in the MLN of WT (n=9), LXRα^−/−^ (*n*=4) and LXRβ^−/−^ (*n*=6) mice. **(f-g)** Representative dot plots and frequencies of Ki-67^+^ **(f)** and AnnexinV^+^ **(g)** Th17 cells in the MLN of WT (*n*=11) and LXRα^−/−^ (*n*=8) mice. Data are represented as means ± SD. *p<0.05, **p<0.01, ***p<0.001, ****p<0.0001 by unpaired Student’s t test or One-Way ANOVA with Bonferroni’s post-test when more than 2 groups.

### LXR signaling in DCs is a prerequisite to restrain RORγt^+^ Tregs in the MLN

Next, we investigated whether LXR signaling was required on T cells to control Th17 generation in the MLN. First, we evaluated if CD4^+^ T cells express LXRα and LXRβ coding transcripts *Nr1h3* and *Nr1h2*, respectively. In line with a previous report on human CD4^+^ T cells ^12^, both isoforms were expressed in CD4^+^ T cells from the MLN, with *Nr1h2* (LXRβ) expression being higher compared to *Nr1h3* (LXRα) (**Supplementary Fig. 1a and 1b**). Next, we investigated whether LXR signaling modulates T cell differentiation *in vitro*. Towards this, splenic CD4^+^ T cells were purified and differentiated *in vitro* under Th2, Treg or Th17 polarizing conditions in presence or absence of LXR agonist (GW3965) or antagonist (GSK2033). While LXR inhibition did not affect CD4^+^ T cell differentiation towards any of the tested subsets, LXR activation reduced differentiation to all the tested Th subsets (**Supplementary Fig. 2a and 2b)**. Further, LXR activation inhibited CD4^+^ T cell proliferation (**Supplementary Fig. 2c and 2d**), in accordance with the previously reported antiproliferative effect of LXR ^17^, suggesting that defects in Th subset differentiation may be due to impaired proliferation. Next, we investigated whether LXR activation in T cells was required to promote Th17 generation *in vivo* using mixed bone marrow (BM) chimera. Given that lack of LXRα results in expansion of only Th17 but not RORγt^+^ Tregs, we focused on LXRα-deficient mice to further understand its role in MLN Th17 cells. We used WT calibrator BM cells from CD90.1 CD45.2 mice that were mixed in 1:1 ratio with BM cells from LXRα^−/−^ or WT mice (characterized by expression of the congenic markers CD90.2 and CD45.2). Mixed BM cells were intravenously injected into WT CD45.1 lethally irradiated mice generating WT:LXRα^−/−^ and WT:WT mixed BM chimeras respectively (**Fig. 2a**). Six-weeks after reconstitution, most of MLN-resident immune cells were donor-derived (**Supplementary Fig. 3a**). Among donor cells, we were able to distinguish RORγt^+^ CD4^+^ T cells from the calibrator (CD90.1) and WT or LXRα^−/−^ (CD90.2) BM-derived cells (**Supplementary Fig. 3a**). Similar to WT:WT chimeras, we found comparable frequencies of RORγt^+^ CD4^+^ T cells between WT and LXRα^−/−^ (**Fig. 2b and c**), indicating that intrinsic deficiency in LXRα does not affect Th17 cell differentiation in the MLN. To further confirm these results, we generated mice lacking LXR in CD4^+^ T cells (LXRαβ^ΔCD4^) by crossing CD4-cre with LXRαβ^flox/flox^ mice. As expected, LXRαβ^ΔCD4^ did not show any difference in frequencies of Th17 and RORγt^+^ Treg cells compared to their LXRαβ^flox/flox^ littermates (**Fig.2d)**. To further investigate the expansion of Th17 cells in LXR-deficient mice, we focused on dendritic cells (DCs), particularly CD103^+^CD11b^+^ DCs, due to their ability to activate and promote differentiation of Th17 cells ^6^. Analysis of DCs (defined as CD45^+^CD11c^+^CD64^neg^MHC-II^+^) showed a significant increase in DC frequencies in LXRα^−/−^ compared to WT, while total number of DCs were comparable (**Supplementary Fig. 4a**). However, further stratification of DCs using CD103 and CD11b expression resulted in higher numbers of CD103^+^CD11b^+^ DCs, but not CD103^+^ or CD11b^+^ single positive DCs in the MLN of LXRα^−/−^ mice compared to WT (**Supplementary Fig. 4b and 4c**), suggesting that Th17 cell expansion might be controlled by the CD103^+^CD11b^+^ DC population. To further confirm, we generated mice lacking LXR in CD11c^+^ cells (LXRαβ^ΔCD11c^). Surprisingly, we did not observe any difference in Th17 cell frequencies between LXRαβ^ΔCD11c^ mice and controls (**Fig. 2e**). However, a significant increase in RORγt^+^ Treg cells was seen in mice lacking LXR in CD11c^+^ DCs compared to control mice (**Fig. 2e**). These results suggest that while LXR signaling in CD11c^+^ DCs modulate the RORγt^+^ Tregs, LXR signaling neither in T cells nor DCs is required to modulate Th17 cells in the MLN.

**Figure 2.**
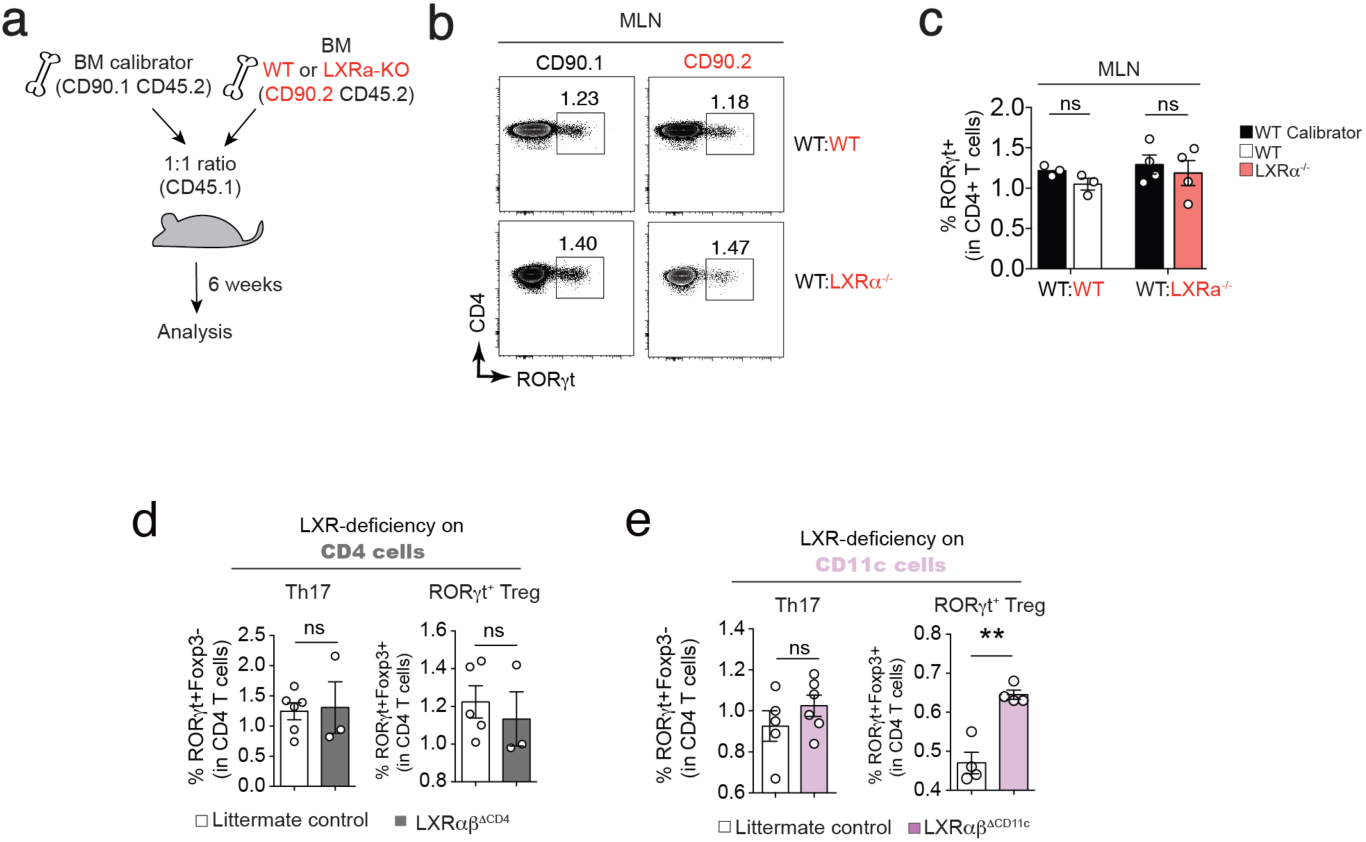
LXR does not intrinsically regulate RORyt^+^ T cells in the MLN. **(a)** Scheme of the mixed bone marrow chimera experiment: bone marrow (BM) cells from WT or LXR□^−/−^ mice (CD90.2 CD45.2) were mixed in a 1:1 ratio with WT calibrator BM cells (CD90.1 CD45.2) and injected into irradiated WT recipients (CD45.1) for the generation of mixed BM chimeras. Six weeks after transfer, MLN from recipient mice were analyzed by flow cytometry. **(b-c)** Representative dot plots (**b**) and frequencies **(c)** of RORyt^+^ cells gated on CD4^+^ T cells from each donor (data are from one representative experiment out of 2 independent experiments). (**d**) Frequencies of Th17 and ROR□t^+^ Tregs gated on CD4^+^ T cells in littermate controls (LXRαβ^flox/flox^) and LXRαβ^ΔCD4^ (CD4-Cre × LXRαβ^flox/flox^) mice. (**e**) Frequencies of Th17 and RORyt^+^ Tregs gated on CD4^+^ T cells in littermate controls (LXRαβ^flox/flox^) and LXRαβ^ΔCD11c^ (CD11c-Cre × LXRαβ^flox/flox^) mice. Each dot represents one mouse and data are represented as means ± SEM. ns=non-significant, *p<0.05, **p<0.01, ***p<0.001, ****p<0.0001 by unpaired Student’s t test.

### LXR signaling is not required in immune or intestinal epithelial cells to regulate Th17 cells in MLN

Mucosal Th17 cell responses have been shown to be dependent on both immune and intestinal epithelial cells (IECs). To further investigate in which compartment LXR activity was needed to influence the MLN Th17 cells and given the ubiquitous expression of LXR we generated mice lacking LXR in IECs (LXRαβ^ΔIEC^) or immune cells (LXRαβ^ΔVAV^) by crossing Villin-Cre and Vav1-iCre with LXRαβ^flox/flox^ mice respectively. Neither LXRαβ^ΔIEC^ nor LXRαβ^ΔVAV^ mice showed any difference in frequencies of Th17 cells compared to their respective littermate controls (**Fig. 3a-d**). On the other hand, as expected LXRαβ^ΔVAV^ mice exhibited a significant increase in frequency of RORγt^+^ Treg cells compared to littermate controls (**Fig. 3c and d**), further confirming our findings with LXRαβ^ΔCD11c^ mice with respect to RORγt^+^ Treg cells. Overall, our data suggest that LXR is not required either in immune or epithelial cells to modulate Th17 cells in the MLN.

**Figure 3.**
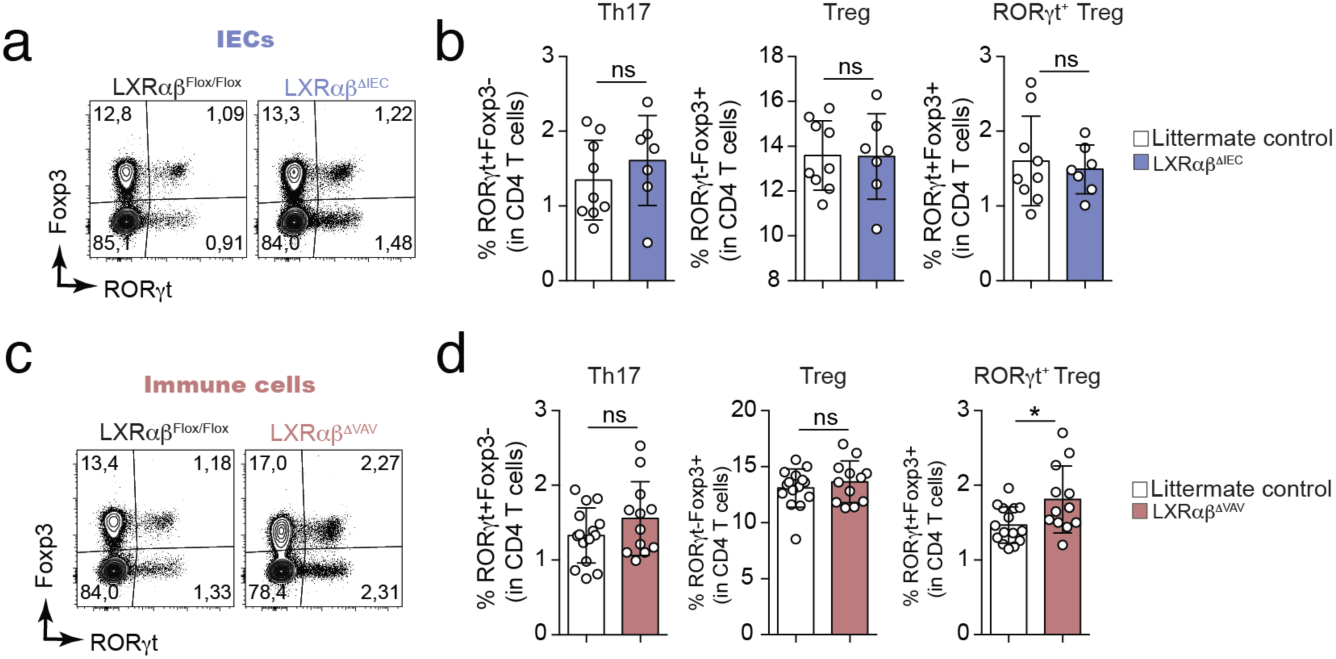
LXR is neither required on immune nor intestinal epithelial cells to regulate MLN Th17. **(a-b)** Representative dot plot **(a)** and frequencies (**b**) of Th17 (RORyt^+^Foxp3^−^), Treg (RORyt^−^Foxp3^+^) and RORyt^+^ Treg (RORyt^+^Foxp3^+^) cells in littermate control (LXRαβ^flox/flox^, n=9) and LXRαβ^ΔIEC^ (Villin-Cre × LXRαβ^flox/flox^, n=7) mice. (**c-d**) Representative dot plot **(c)** and frequencies (**d**) of Th17 (RORyt^+^Foxp3^−^), Treg (RORyt^−^Foxp3^+^) and RORyt^+^ Treg (RORyt^+^Foxp3^+^) cells in littermate control (LXRαβ^flox/flox^, n=15) and LXRαβ^ΔVAV^ (Vav1-iCre × LXRαβ^flox/flox^, n=12) mice. Data are represented as means ± SD. ns=non-significant, *p<0.05, **p<0.01, ***p<0.001, ****p<0.0001 by unpaired Student’s t test.

### Horizontal microbiota transfer from LXRα deficient mice partly restores MLN Th17 cells in WT

The microbiota is considered one of the major drivers of Th17 cell generation in the intestine, as seen by virtually complete absence of Th17 cells in germ-free mice ^10^. As the lack of LXR in either immune or IEC compartment did not explain the altered Th17 cell frequency seen in LXRα^−/−^ mice, we reasoned that alternative factors, such as the microbiota, might cause the observed phenotype. To address the bacterial contribution to MLN Th17 cell generation in WT and LXRα^−/−^ mice by horizontal and/or vertical transfer of bacteria to suckling or adult mice, we performed cross-fostering and co-housing experiments. To evaluate whether bacterial shaping early in life during the lactating period could affect MLN Th17 cells in adulthood, we performed a cross-fostering experiment where newborn WT pups were cross-fostered at birth with LXRα^−/−^ dams and vice versa (**Fig. 4a**). Four weeks after birth, mice were weaned into separate cages based on their genotypes and MLN T cell composition was analyzed around 8-12 weeks after birth (**Fig. 4a**). WT mice displayed decreased frequencies of Th17 cells in the MLN compared to LXRα^−/−^ mice regardless of fostering conditions, thus suggesting that the microbiota or nutrients transferred vertically by breastfeeding was not causative of the observed expansion of Th17 cells in LXRα^−/−^ mice (**Fig. 4b**). Next, to investigate whether horizontal transmission of the microbiota during adulthood might account for Th17 expansion, WT and LXRα^−/−^ non-littermate mice from the same provider were either co-housed or single housed according to their genotype for 4 weeks (**Fig. 4c**) to allow for normalization of microbiota ^18^. Of note, WT mice co-housed with LXRα^−/−^ mice partially gained higher frequencies of MLN Th17 cells compared to single housed WT mice. However, similar to what was observed in cross fostered mice, Th17 cells were proportionally higher in the MLN of LXRα^−/−^ mice compared to WT regardless of housing conditions (**Fig. 4d and e**). Altogether, these data suggest that a distinctive horizontally transmitted gut microbiota in LXRα^−/−^ mice partly induce the expansion of Th17 cells in the MLN.

**Figure 4.**
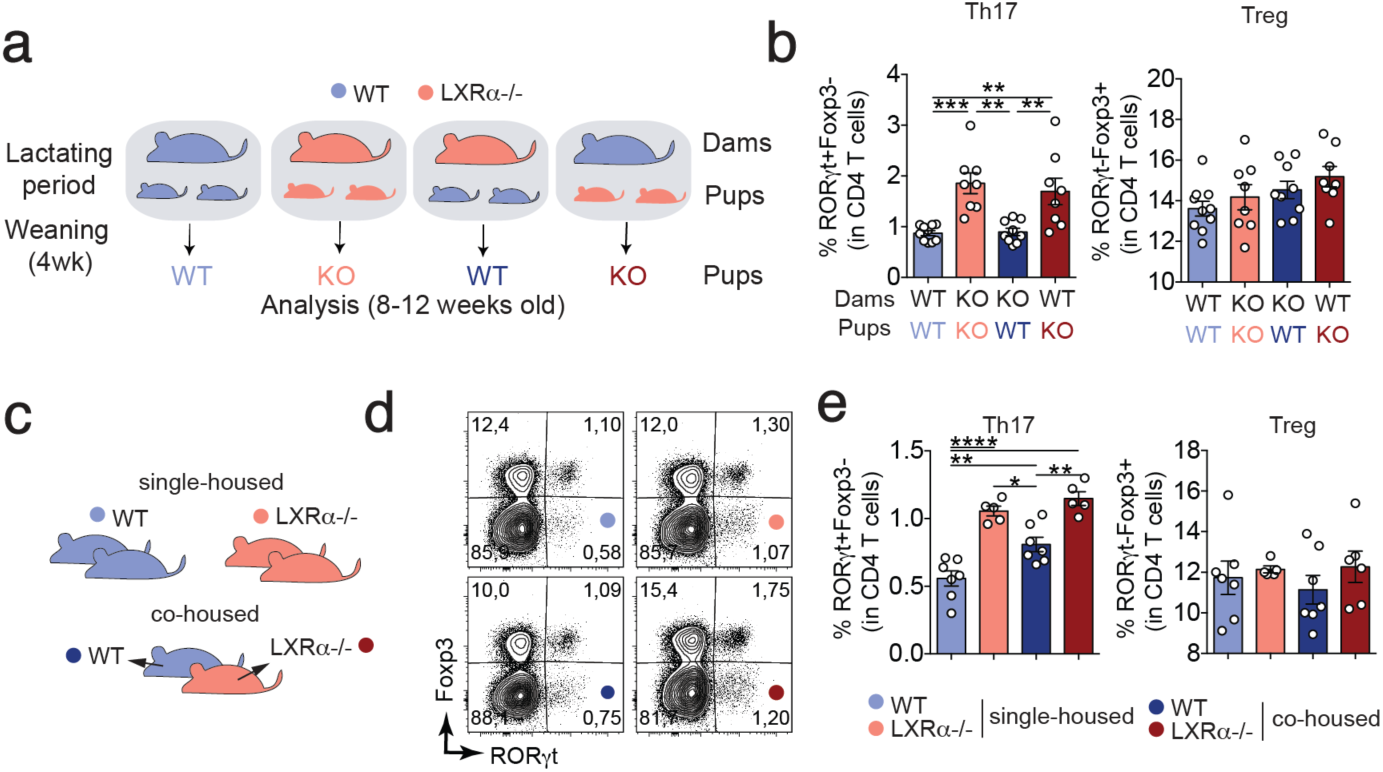
Horizontal microbiota transfer from LXR deficient mice only partially increases MLN Th17 in WT mice. **(a)** Schematic representation of the cross-fostering experiment: WT and LXRα^−/−^ newborn pups were either left with their biological mothers or they were swapped between their mothers (foster mother) right after birth and for the entire lactating period. Four weeks after birth, mice were weaned and kept in separate cages based on genotype and fostering mother. Mice were then sacrificed at 8-12 weeks of age and MLN was analyzed by flow cytometry. **(b)** Frequencies of Th17 and Treg cells out of CD4 T cells in the MLN of WT and LXRα^−/−^ mice stratified based on genotype and foster mother (each dot represents one mouse out of 2 independent experiments). **(c)** Schematic representation of the co-housing experiment: WT and LXRα^−/−^ mice coming from different colonies were either housed separately (single-housed) based on their genotype or together (co-housed) in the same cage for a period of 4 weeks. Representative dot plots **(d)** and quantification of frequencies **(e)** of Th17 and Treg cells in the MLN of WT and LXRα^−/−^ mice either single- or co-housed (each dot represents one mouse out of 4 independent experiments). Data are represented as means ± SEM. ns=non-significant, *p<0.05, **p<0.01, ***p<0.001, ****p<0.0001 by one-way ANOVA with Bonferroni’s post-test.

### LXRα mediated regulation of MLN Th17 is independent of antibiotic-sensitive intestinal microbiota

Since the presence of SFB is sufficient to induce Th17 cell priming shortly after colonization ^10,19^, we evaluated if SFB levels correlate with Th17 frequencies in WT and LXRα^−/−^ mice. Increased frequency of Th17 cells in the MLN of LXRα^−/−^ mice did not correlate with SFB abundance (**Fig. 5a**), suggesting that Th17 expansion in LXRα-deficient mice is SFB-independent. We, therefore, tested if other bacterial communities were different between WT and LXRα^−/−^ mice and thus potentially explain the expansion of MLN Th17. Towards this we performed 16S bacterial rRNA sequencing of WT and LXRα^−/−^ littermate mice (**Fig. 5b**). Of note, we observed significantly higher enrichment of bacterial species belonging to the family Ruminococcaceae and Lachnospiraceae in LXRα^−/−^ mice compared to WT littermates (**Fig. 5b**), indicating that LXRα modulates the composition of the intestinal microbiota. To further test if the distinct microbiota composition accounts for the differences in MLN Th17 cells, we treated WT and LXRα^−/−^ littermates for 10 days with a broad-spectrum antibiotic cocktail. As expected, we observed a dramatic change in the microbiota composition, which was dominated by Enterobacteriaceae in both WT and LXRα^−/−^ mice (**Fig. 5c)**. Interestingly, we detected the presence of opportunistic bacteria, such as Bacillaceae and Listeriaceae only in the LXRα-deficient mice regardless of the treatment or not with antibiotics (**Fig. 5d**). However, as expected antibiotics treatment resulted in significant abrogation of majority of bacteria in the intestinal tract (**Fig. 5e**), including SFB (**Fig. 5f**), Ruminococcaceae and Lachnospiraceae (**Fig. 5g**). Having demonstrated that the antibiotic treatment depletes majority of the intestinal microbiota in the LXRα-deficient background, we analyzed the levels of MLN Th17 cells. While the antibiotics treatment resulted in reduction of MLN Th17 cells in WT mice, it did not alter Th17 frequencies in LXRα^−/−^ mice (**Fig. 5h**), suggesting the expansion of MLN Th17 in LXRα^−/−^ mice is independent of antibiotic-sensitive microbiota. Whether the presence of antibiotic-resistant bacteria is sufficient to induce MLN Th17 cells in LXRα^−/−^ mice further remains to be addressed. Taken together, our results demonstrate that LXRα extrinsically regulates MLN Th17 expansion in a SFB-independent manner.

**Figure 5.**
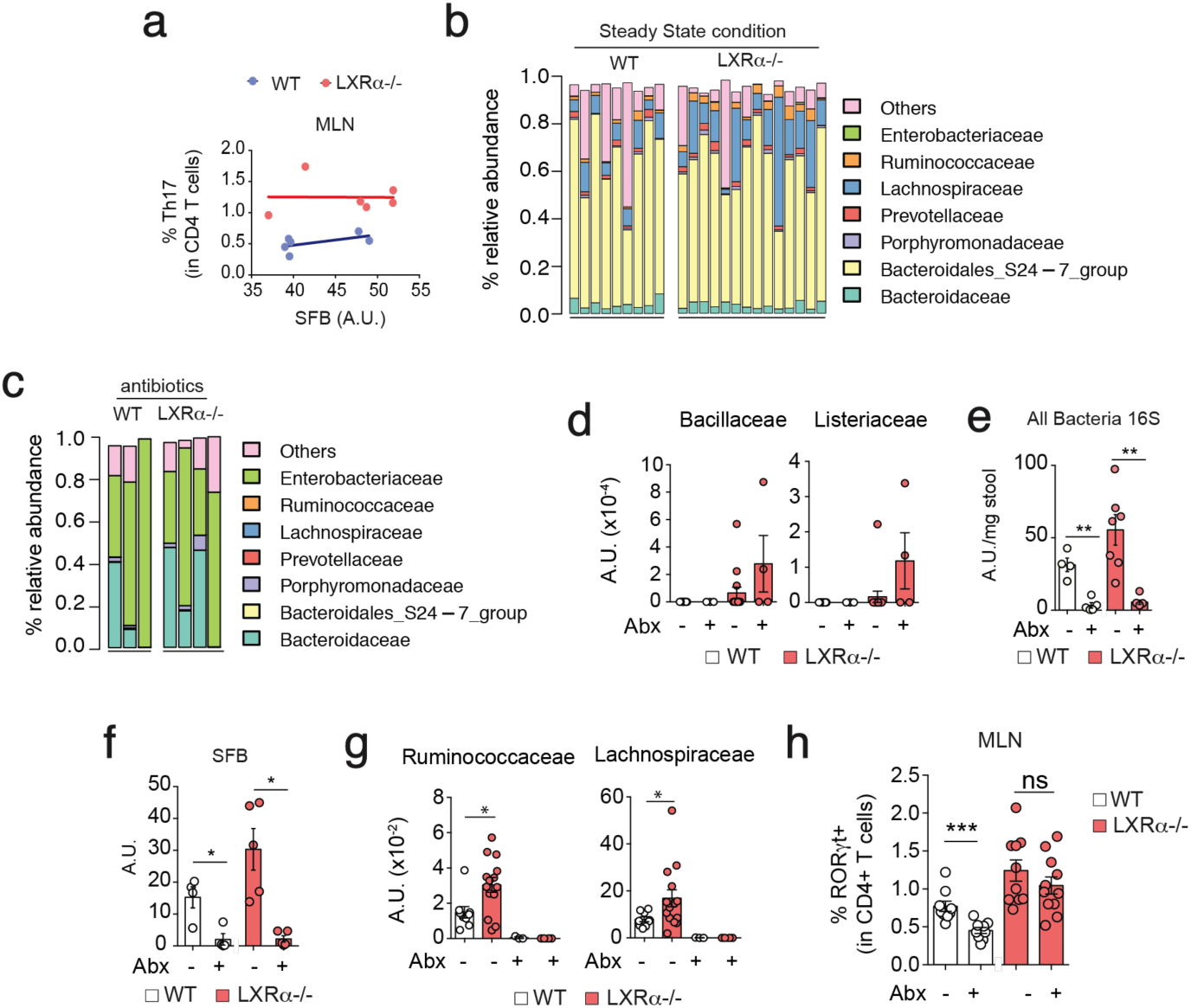
LXRα dampens MLN Th17 expansion in a SFB-independent manner. **(a)** Relative amount of total bacteria (quantified by universal 16S) and SFB in WT and LXRα^−/−^ mice. Universal bacteria units were normalized by mg of stool whereas SFB units were normalized by universal bacteria values **(b)** Correlation between Th17 frequencies in the MLN and SFB relative amount in the stool samples from WT and LXRα^−/−^ mice. **(c-d)** Relative abundance of indicated bacterial families in the stool samples from WT and LXRα^−/−^ mice in steady state (**c**) and after antibiotics (Abx) treatment (**d**) as determined by 16S rRNA sequencing. (**e**) Plot showing normalized count of Bacillaceae and Listeriaceae families obtained from 16S rRNA sequencing in WT and LXRα^−/−^ mice before and after Abx treatment. (**f**) Relative amount of total bacteria per mg of stool in WT and LXRα^−/−^ mice before and after Abx treatment. (**g**) Relative amount of SFB (normalized to universal 16S) in WT and LXRα^−/−^ mice before and after Abx treatment. (**h**) Plot showing normalized count of Ruminococcaceae and Lachnospiraceae families obtained from 16S rRNA sequencing in WT and LXRα^−/−^ mice before and after Abx treatment. (**i**) Frequencies of Th17 in the MLN of WT (n=9 untreated, n=9 Abx treated) and LXRα^−/−^ (n=10 untreated, n=11 Abx treated) mice treated or not with antibiotics. Each dot represents one mouse and data are represented as means ± SEM. ns=non-significant, *p<0.05, **p<0.01, ***p<0.001, ****p<0.0001 by unpaired Student’s t test.

## Discussion

LXR has been shown to play an anti-inflammatory role by dampening the differentiation of Th17 cells in a T cell intrinsic manner. Here, using mice lacking LXRα, LXRβ or both isoforms in different cell compartments, we have defined a model in which LXR signaling controls the generation of two different T helper cells by distinct mechanisms. While LXRα extrinsically restrains expansion of Th17 cells, at least in part by horizontally transferred environmental factors, RORγt^+^ Treg cells instead were controlled by LXR signaling in CD11c^+^ myeloid cells.

While Th17 cells are important mediators of antibacterial/fungal immunity, they also play a key role in the pathogenesis of autoimmune/inflammatory diseases. In agreement with the high density of bacteria colonizing the gut mucosa of microbiota, the intestine harbors the majority of Th17 cells present in our body. Gut-draining lymphoid tissues, including MLN, are the main sites orchestrating the priming of adaptive immune responses to intestinal luminal antigens. Here, we dissected the relative contribution of each LXR isoform and found that while lack of LXRα results in a selective increase only in Th17, lack of LXRβ controls all RORγt-expressing CD4^+^ T cells, including Th17 and RORγt^+^ Tregs. Previous studies have claimed that LXR-mediated control of Th17 differentiation occurs in a T cell-intrinsic fashion, as shown by *in vitro* T cell differentiation assays ^15,17^. However, by using two different *in vivo* approaches (mixed BM chimera and conditional depletion of LXR in CD4^+^ T cells), we showed that LXR is not required on T cells to control Th17 differentiation in the MLN. A recent study reported that LXR deficiency and cholesterol accumulation in dendritic cells altered their antigen presenting function, thus affecting priming and expansion of adaptive immune cells ^20^. Our analysis of MLN DC subsets in mice lacking LXRα, showed higher numbers of CD103^+^CD11b^+^ DCs, previously implicated in the induction of intestinal Th17 differentiation ^6^. Nevertheless, our results show that conditional ablation of LXR signaling in CD11c^+^ myeloid cells (or in total immune cells) did not result in any alteration in MLN Th17 cells. By contrast, mice lacking LXR in CD11c^+^ myeloid cells resulted in increased RORγt^+^ Tregs in the MLN. These findings pose several appealing questions that remain to be answered: is the generation of RORγt^+^ Tregs dependent on CD103^+^CD11b^+^ DCs as shown for Th17 ^6^? Does altered cholesterol metabolism in DCs skew their potential towards the preferential generation of RORγt^+^ Treg? Are RORγt^+^ Tregs generated by LXR-deficient DCs functionally comparable to the ones generated by LXR-sufficient DCs? While these questions await experimental validation, our results thus far suggest that LXR signaling is necessary in antigen-presenting cells to restrain the expansion of RORγt^+^ Treg cells.

The intestinal epithelium is the first cellular line detecting environmental cues in the lumen and transmitting those signals to underlying immune cells in the lamina propria. Due to these properties, IECs have been implicated in shaping the differentiation and activity of lymphocytes, including Th17 cells ^10,21^. However, selective LXR deficiency in IECs alone was not sufficient to expand MLN Th17 cells unlike the whole body LXR deficient mice. These findings could potentially be explained by the requirement of LXR either in non-immune/non-epithelial cells or in immune and epithelial cells simultaneously to control MLN Th17.

With the aim of investigating what contributes to the specific expansion of MLN Th17 in LXRα^−/−^ mice, we tested the relative contribution of the microbiota. Cross-fostering and co-housing experiments suggested that only horizontal transfer of bacteria in adulthood was sufficient to partially increase Th17 cells in WT mice, although this did not reach the level of Th17 cells observed in the MLN of LXRα^−/−^ mice. These results propose a model whereby a combination of genetic deficiency of LXR and microbiota composition controls MLN Th17. We thus used littermate and cohoused mice to further reduce any external or LXR-independent modulation of the microbiota. In these experiments, we observed significantly higher enrichment of bacteria belonging to the family Ruminococcaceae and Lachnospiraceae in LXRα^−/−^ mice compared to WT littermates in steady state, suggesting that lack of LXRα allows preferential colonization of specific groups of bacteria. Both Ruminococcaceae and Lachnospiraceae belong to the order Clostridiales and phylum Firmicutes and have been associated with multiple human diseases including IBD ^22-24^ and atherosclerosis ^25,26^. While the abundance of both Ruminococcaceae and Lachnospiraceae are decreased in IBD patients, suggesting a protective role, they have been shown to be associated with pro-atherogenic effects. Since Th17 cells have been shown to play context dependent functions in both IBD ^27,28^ and atherosclerosis ^29^, future studies will address whether LXRα mediated regulation/restraint of Ruminococcaceae and Lachnospiraceae impact the diseases such as IBD and atherosclerosis in a Th17 cell-(in)dependent manner.

Segmented filamentous bacteria (SFB) are sufficient to induce Th17 cells in the intestine ^10^. However, in our microbiota analysis we did not observe any difference in the microbiota composition belonging to the family Clostridiaceae (to which SFB belongs) between WT and LXRα^−/−^ mice. Furthemore, we did not observe any difference in the amount of SFB in the colonic stool of WT and LXRα^−/−^ mice as well as no correlation between the Th17 cells in the MLN with the amount of SFB in the stool samples. While the lymph node that drains the intestine, i.e. MLN, serves as the site of T effector cell priming by antigen presenting cells such as DCs, Goto *et al*. ^30^ demonstrated that priming of SFB-induced Th17 happens in the intestinal lamina propria rather than the draining lymph nodes. Therefore, our observation of SFB-independent increase of Th17 in the MLN of LXRα^−/−^ mice is supported by the notion that SFB-mediated Th17 priming happens *in loco* in the intestinal lamina propria without affecting the Th17 cells observed in the MLN ^30^. To further confirm, if the increased Th17 observed in LXRα^−/−^ mice was due to any other bacterial species, we treated both WT and LXRα^−/−^ mice with broad-spectrum antibiotics. While there was a reduction in total bacteria in both WT and LXRα^−/−^ mice, we observed a reduction of MLN Th17 only in WT mice. Moreover, 16S rDNA sequencing showed no significant differences in microbiota composition between WT and LXRα^−/−^ mice after antibiotic treatment. Interestingly, we occasionally detected the presence of the opportunistic bacteria, such as Bacillaceae and Listeriaceae only in the LXRα-deficient mice with or without antibiotics treatment. Whether, the presence of such opportunistic bacteria influences the MLN Th17 cells in LXRα^−/−^ mice needs further investigation. These findings suggest that while microbiota influences the priming of Th17 in the MLN of WT mice, in LXRα^−/−^ mice it is microbiota-independent. Although bacterial species constitute the bulk of intestinal microbiota, we cannot rule out if the observed increase in MLN Th17 cells in LXRα^−/−^ are dependent on other minor constituents of the microbiota such as antibiotic resistant and opportunistic bacterial species, fungi, viruses and protists etc. Further studies are necessary to address these questions.

Overall, using multiple approaches and genetic tools, we demonstrate that LXR restrains the expansion of T helper subsets such as Th17 and RORγt^+^ Tregs by distinct mechanisms in the MLN. Contrary to previous findings ^15^, we demonstrated that the effect of LXR on Th17 is T cell extrinsic and that LXRα isoform is sufficient to restrain only Th17 and not RORγt^+^ Tregs. We further demonstrate that LXR is required neither on immune nor IECs and is independent of antibiotic-sensitive intestinal bacteria, thus suggesting a new mode of regulation of MLN Th17 that might rely on other untested cell types (e.g. stromal or neuronal cells etc.) or a simultaneous requirement on multiple cell types (e.g. immune and epithelial together). On the other hand, RORγt^+^ Tregs were dependent on LXR signaling on CD11c^+^ myeloid cells and was specifically modulated by LXRβ isoform. This demonstrates LXR isoform-specific modulation of Th subsets, which can be further explored to understand the functional relevance of these Th subsets both in intestinal and extra intestinal tissues under different disease conditions such as IBD, EAE, SLE, atherosclerosis etc.

## Materials and Methods

### Mice

Mice, aged 6-20 weeks of C57BL/6J background were used for all experiments. Wildtype (CD45.2 and CD90.2), CD45.1 and CD90.1 were purchased from either Taconic (Taconic, Ry, Denmark) or from mice bred at the Microbiology, Tumor and Cell Biology (MTC), Karolinska Institutet (CD45.1 mice). CD11c-cre, CD4-cre and Vav1-icre mice were purchased from Jackson Laboratories and bred locally. LXRα^−/−^, LXRβ^−/−^, Villin-cre and LXRαβ^flox/flox^ mice were kindly provided by professor Jan-Åke Gustafsson (Karolinska Institutet, Huddinge). Animals were maintained under specific pathogen-free conditions at the AFL or MTC animal facility and handled according to protocols approved by the Institutional Animal Care and Use Committee at the Karolinska Institutet (Stockholm, Sweden). For LXR activation *in vivo*, mice were fed with either a control diet or diet formulated with GW3965 (50mg/kg/day) with AIN 93G diet as the basal diet (SSNIFF, Germany).

### Single cell suspensions and *in vitro* cultures

MLN cells were isolated by smashing the lymph nodes through a 70 µm cell strainer. Red blood cells (spleen) were lysed using ACK buffer. Remaining cells were pelleted for further use. For *in vitro* T-cell differentiation, splenic CD4^+^ T cells were enriched by positive selection using the CD4 (L3T4) isolation kit (Miltenyi) following the manufacturer’s instructions. For polarizing experiments CD4^+^ T cells were stimulated with 1µg/mL soluble anti-CD3/CD28 in the presence or absence of 5 µg/mL GSK 2033 or GW3965 and incubated at 37°C with 5% CO_2_ for 5 days. T cell subsets were analyzed by FACS on day5. Following polarizing conditions were used: 2 ng/mL IL-2 (Th0); 2 ng/mL IL-2, 30 ng/mL IL-4, and 1.25 μg/mL anti-IFN-γ (Th2); 2 ng/mL IL-2, 3 ng/mL TGF-β, 30 ng/mL IL-6, 30 ng/mL IL-23, 0.625 μg/mL anti-IL-4 and 1.25 μg/mL anti-IFN-γ (Th17); 2 ng/mL IL-2 and 2 ng/mL TGF-β (Treg).

### Co-housing

3-4 weeks after birth, mice coming from separate homozygous colonies (either WT or LXRa^−/−^) were weaned based on sex. From week 6 after birth until sacrifice mice were then either left single-housed (i.e. housed with littermates with the same genotype) or co-housed with mice of the opposite genotype for 4 weeks until sacrifice.

### Cross-fostering

Breeding cages with 2 females and 1 male mouse with the same genotype (i.e. either WT × WT or LXRα^−/−^ × LXRα^−/−^) were paired. After three weeks, pregnant females were separated and single-housed in new cages. The following week, when pups were born, the litters were either kept in the same cage or swapped and added to cages with foster female of the opposite genotype. Four weeks after birth, pups were weaned into separate cages based on genotype and sex. Mice were then sacrificed for analysis when 10-19 weeks old.

### Antibiotics treatment

WT and LXRα^−/−^ mice were treated with antibiotic cocktail for 10 consecutive days. The following cocktail was given by oral gavage: Ampicillin (1mg/ml), Kanamycin (1mg/ml), Gentamicin (1mg/ml), Metronidazole (1mg/ml), Neomycin (1mg/ml), and Vancomycin (0.5mg/ml). After antibiotic treatment, mice were sacrificed, and organs and stool samples were collected for subsequent analysis.

### Statistics Analysis

Statistical analyses were performed with GraphPad Prism version 4.01 (GraphPad Software, Inc., 2005). Immune responses among groups of mice are presented as means with standard errors. Comparisons of mean immune responses were performed using two-sided t-tests. In all cases, p-values of less than 0.05 were considered significant.

## Acknowledgments

We thank members of the Villablanca lab for helpful comments. SD was supported by grants from Cancerfonden (CAN 2016/1206). AF was supported by a grant from the German research association (DFG, 808021). EJV was supported by grants from the Swedish Research Council VR grant K2015-68X-22765-01-6, FORMAS grant nr. FR-2016/0005 and Wallenberg Academy Fellow (WAF) program.

## Author contributions

S.M.P., S.D., A.F., R.F.C. and C.D. designed and performed most experiments, analyzed the data, and wrote the manuscript. K.P.J and Y.H. performed analysis. P.A., and J.A.G. provided mice and feedback. E.J.V. conceived the study, designed experiments, analyzed the data, and wrote the manuscript.

