## Supplementary text and Figures for "Liver X Receptor regulates Th17 and RORγt^+^ Treg cells by distinct mechanisms"

#### Supplementary Methods

##### Mixed BM chimera

Recipient CD45.1 wildtype mice were irradiated (6 Gy) twice, 3 hours apart. Bone marrow (BM) from donor mice was isolated by mechanical disruption of bones and filtered. 5 million WT calibrator BM cells (CD45.2, CD90.1) were mixed in a 1:1 ratio with WT or LXRα$^{-/-}$ BM (all CD45.2, CD90.2) in PBS and transferred intravenously into irradiated CD45.1 recipient mice. Mice were then administered Neomycin sulfate (2mg/mL) via drinking water for 2 weeks. Recipient mice were analyzed by flow cytometry 9 -12 weeks after BM transfer.

##### Flow cytometry

Single cell suspensions were incubated for 10 min at 4°C with Fc-blocking (CD16/32) antibody (eBioscience) and fixable viability dye eFluor780 (eBioscience) prior to staining with fluorochrome-conjugated antibodies for 15 min at 4°C. For staining of intracellular transcription factors, after cell surface staining, cells were incubated with Fixation and Permeabilization buffer (Foxp3 staining kit, eBioscience) for 30 min at 4°C followed by staining for 20 min at room temperature with fluorochrome-conjugated antibodies against transcription factors. The following antibodies were purchased from eBioscience: CD4-PECy7 (RM4-5), CD90.1-FITC (His51), Foxp3-APC (FJK-16S), GATA3-PE (TWAJ), Ki67-PE (20Raj1), CD103-PE (2E7). The following antibodies were purchased from Biolegend: CD4-PerCP (GK1.5), CD11c-APC (N418), CD3-BV785/BV421 (145-2c11), CD45.2-BV650 (104) and CD45.1-PB (A20), MHCII-PerCP (M5/114.15.2), CD64-PB (54-5/7.1). The following antibodies were purchased from BD Biosciences: CD11b-BV785 (M1/70), CD90.2-PECy7 (53-2.1) and RORγt-PE-TR (Q31-378), AnnexinV-FITC. DAPI or Live/Dead Fixable viability dyes (eBioscience) were used to exclude dead cells. All the experiments were acquired using FACS Canto II or FACS LSR Fortessa flow cytometers (BD Biosciences) and analyzed with FlowJo software (TreeStar).

##### RNA extraction, cDNA synthesis and RT-qPCR

Cells were lysed and RNA was extracted using the RNeasy Plus Mini Kit from Qiagen according to the manufacturer's instructions. cDNA was synthesized using iScript cDNA synthesis kit from Bio-Rad according to the manufacturer's instructions. A 10-µl real-time PCR reaction included 2-4 µl cDNA, 5 µl iTaq™ Universal SYBR® Green Supermix (Bio-Rad), primers and water as indicated by the manufacturer's instructions. qPCR was carried out for 40 cycles with 95 °C melting (30 s), 58 °C annealing (30 s), and 72 °C extension (30 s). All reactions were performed on a CFX384 Touch Real-Time PCR. Primer sequences were as follows:

LXRα Forward: 5′- GTTTCTCCTGATTCTGCAACG-3′,

LXRα Reverse: 5′-TCCAACCCTATCCCTAAAGCAA-3′;

LXRβ Forward: 5′- CACCATTGAGATCATGTTGC-3′,

LXRβ Reverse: 5′- GAACTCGAAGATGGGATTGATG-3′;

Hprt Forward: 5′-TCAGTCAACGGGGGACATAAA -3′,

Hprt Reverse: 5′-GGGGCTGTACTGCTTAACAGC-3′;

**DNA extraction from stool and microbiota analysis by 16S sequencing and qPCR**

Stool samples collected at indicated time points were stored at -80 °C until genomic DNA extraction. Bacterial DNA was extracted and purified from fecal samples using either QIAamp Fast DNA Stool Mini Kit (Qiagen) or Maxwell 16 Tissue DNA Purification Kit (Promega) following the manufacturer's instructions. For 16S sequencing, the V3/V4 hypervariable regions from 16S rRNA genes were amplified with primers 341F - 805R using the 1-step PCR procedure as previously described by (*32*). The indexed 16S sequencing libraries of each sample together with PCR controls were pooled at equimolar amount and subjected to Illumina MiSeq sequencer (Illumina, San Diego) using kit v3. After conducting base-calling and demultiplexing, paired-end reads from each sample were generated and subsequently sent for sequencing quality control to guarantee the laboratory performance and sequencing depth. To generate error-free amplicon sequence variants (ASVs), we conducted DADA2 denoise program (v. 1.12) on the quality-filtered reads in R (www.R-project.org) following DADA2 pipeline(*33*). The primer regions on the raw reads were first trimmed off, and then the reads were quality-filtered with DADA2's default setting before subjected to the denoise program. The denoised paired-end reads were then merged with at least 30 bases overlap to recover ASVs. At last, the ASVs were annotated with SILVA database (v.128) (*34*) using DADA2 assign Taxonomy function. Real time qPCR analysis was performed by using universal or SFB specific primers for 16S rRNA genes. A 10-μl real-time PCR reaction included 2-4 μl bacterial DNA (40 ng of bacterial DNA), 5 μl iTaq™ Universal SYBR® Green Supermix (Bio-Rad), primers and water as indicated by the manufacturer's instructions. qPCR was carried out for 40 cycles with 95 °C melting (30 s), 58 °C (for SFB) or 63 °C (All Bact) annealing (30 s), and 72 °C extension (30 s) as described above.

Primers used for library preparation for 16S sequencing:

16S (341F) Forward: 5'-CAAGCAGAAGACGGCATACGAGAT-N8-GTCTCGTGGGCTCGGAGATGTGTATAAGAGACAGGACTACHVGGGTATCTAATCC-3';

16S (805R) Reverse: 5´AATGATACGGCGACCACCGAGATC-N8-TCGTCGGCAGCGTCAGATGTGTATAAGAGACAGCCTACGGGNGGCWGCAG-3′;

Primers used for bacterial qPCR:
16S (All bacteria, UniF340) Forward: 5′- ACTCCTACGGGAGGCAGCAGT -3′,
16S (Total bacteria, UniR514) Reverse: 5′- ATTACCGCGGCTGCTGGC-3′;
SFB (SFB736F) Forward: 5'- GAC GCT GAG GCA TGA GAG CAT-3'
SFB (SFB844R) Reverse: 5'- GAC GGC ACG GAT TGT TAT TCA-3'

For the correlation analysis of SFB levels with MLN Th17 cells, the A.U. of SFB (inverse ΔCT) was calculated as SFB (A.U.) = 1/ΔCT, where ΔCT = Ct for SFB – Ct for All Bacteria.

To calculate the bacterial load, 40 µl real-time PCR reaction included 100ng DNA, 1 µl 16S primers, 10 µl iTaq™ Universal SYBR® Green Supermix (Bio-Rad) and water were used. Ct values were normalized by mg of feces. To calculate SFB A.U. Ct values for SFB were normalized to universal 16S quantification and relative quantification were calculated by 2^-(ΔΔCt) method, using one antibiotic treated mice as calibrator.

#### Supplementary Figure Legends

**Supplementary Figure 1. Both isoforms of LXR are expressed in $CD4^+$ T cells in the MLN.**
RT-PCR **(a)** and qPCR analysis **(b)** of *Nr1h3* (LXRα) and *Nr1h2* (LXRβ) on the indicated FACS sorted cell populations from spleen, MLN, small intestine intra-epithelial compartment (SI IEL) and small intestine lamina propria (SILP).

**Supplementary Figure 2. LXR activation dampens T cell proliferation and differentiation.**
Splenic T cells were differentiated in vitro for 5 days under none (Th0), Th2, Th17 or Treg polarizing conditions in the presence of vehicle (DMSO), 5 µM LXR agonist (GW3965) or 5 µM LXR antagonist (GSK2033). **(a)** Representative dot plots showing RORγt expression on CD4 T cells at the end of the experiment. **(b)** Frequencies of *in vitro* differentiated Th2 ($Gata3^+$), Th17 ($ROR\gamma t^+$) or Treg ($Foxp3^+$) cells out of $CD4^+$ T cells in the presence of vehicle, LXR agonist (GW3965) or antagonist (GSK2033). (**c**) Histogram showing dilution of cell trace violet of *in vitro* differentiated Th17 ($ROR\gamma t^+$) cells in the presence of vehicle, LXR agonist (GW3965) or antagonist (GSK2033). (**d**) Quantification of % cell trace violet diluted cells (indicator of % proliferation) of *in vitro* differentiated Th17 cells in the presence of vehicle, LXR agonist (GW3965) or antagonist (GSK2033).

**Supplementary Figure 3. Gating strategy to analyze the mixed BM chimera experiment.**

(**a**) Scheme showing the gating strategy used to analyze the mixed bone marrow chimera experiment. Donor-derived cells were gated as CD45.2$^+$ cells and different donors were discriminated based on CD90 expression. Frequencies of RORγt$^+$ T cells were analyzed within each donor-derived cells.

**Supplementary Figure 4. LXRα regulates CD103$^+$CD11b$^+$ DCs in the MLN.**

(**a**) Flow cytometry analysis of frequencies and numbers of CD11c$^+$ MHCII$^+$ dendritic cells (DC) in the MLN of WT (*n*=18) and LXRα$^{-/-}$ (*n*=21) mice. **(b-c)** Frequencies **(b)** and absolute numbers (**c**) of CD103$^+$, CD103$^+$CD11b$^+$, CD11b$^+$ DC subsets in the MLN of WT (*n*=18) and LXRα$^{-/-}$ (*n*=21) mice (four independent experiments). Data are represented as means ± SD. ns=non-significant, *p<0.05, **p<0.01, ***p<0.001, ****p<0.0001 by unpaired Student's t test.

### Supplementary Figure 1

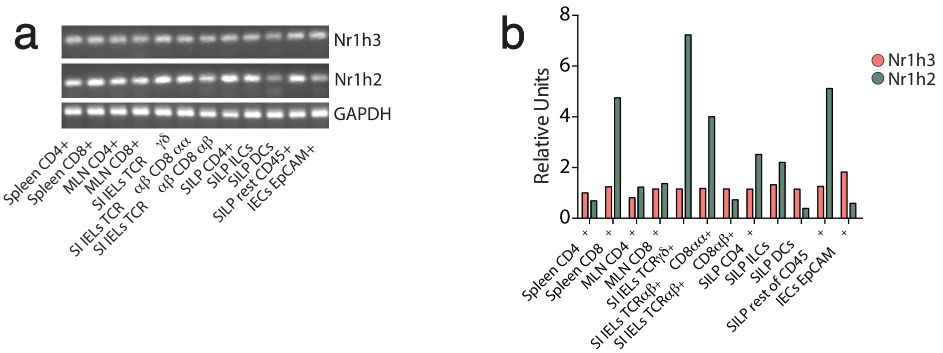

### Supplementary Figure 2

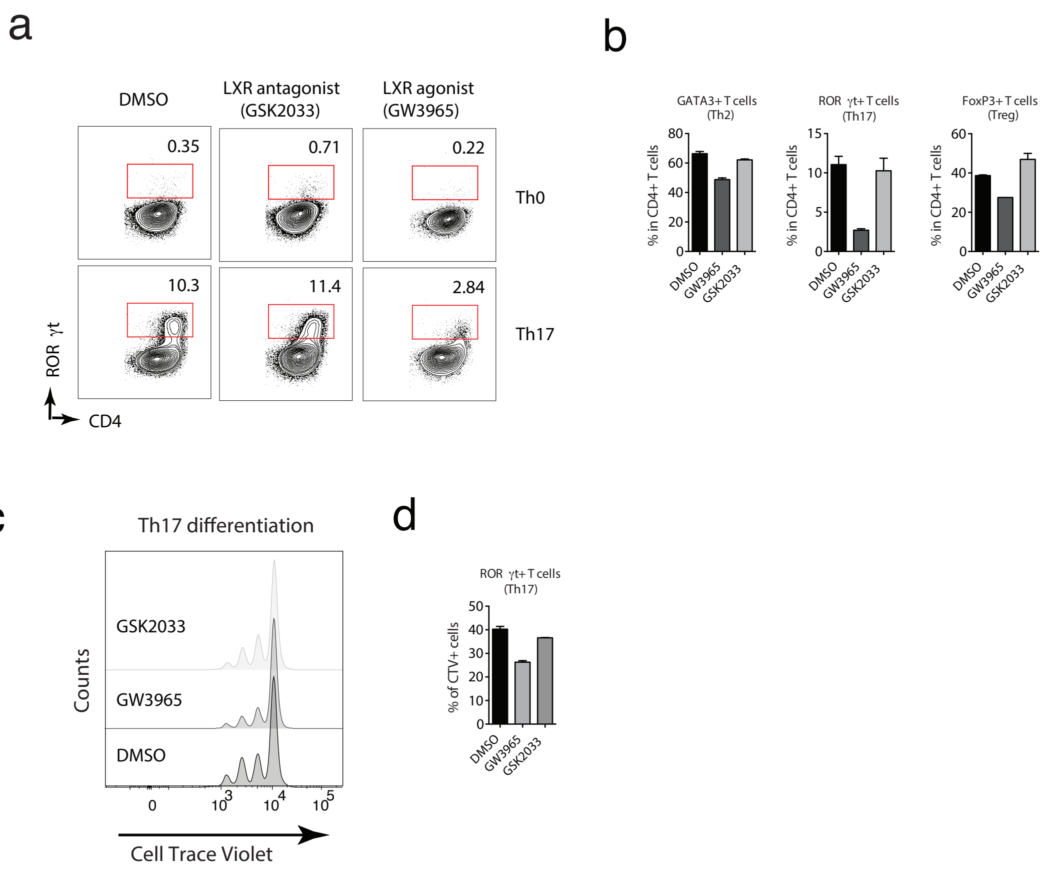

### Supplementary Figure 3

a

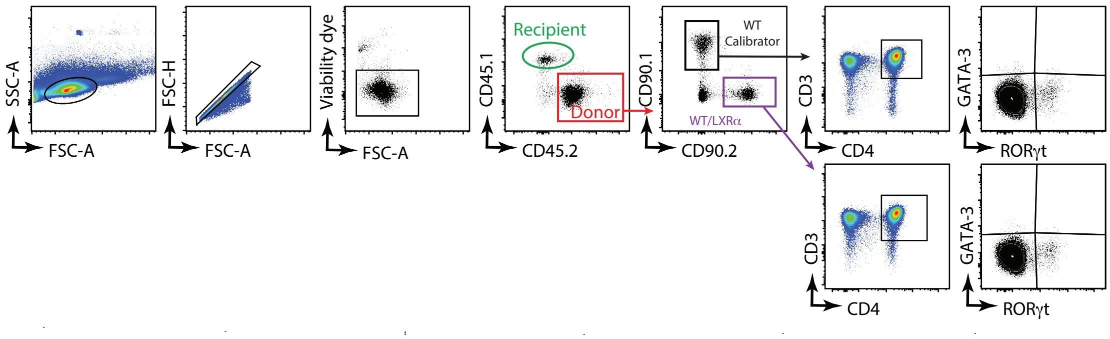

### Supplementary Figure 4

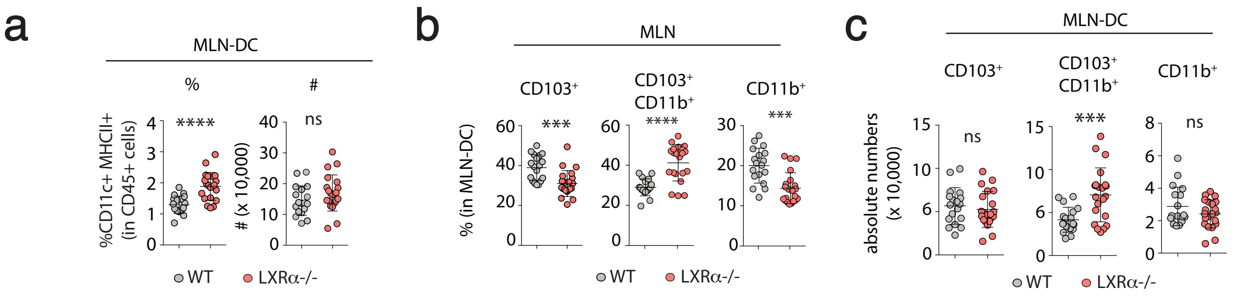
